# The evolution of a super-swarm of foot-and-mouth disease virus in cattle

**DOI:** 10.1101/512178

**Authors:** Jonathan Arzt, Ian Fish, Steven J. Pauszek, Shannon L. Johnson, Patrick S. Chain, Devendra K. Rai, Elizabeth Rieder, Tony L. Goldberg, Luis L. Rodriguez, Carolina Stenfeldt

**Affiliations:** Foreign Animal Disease Research Unit, Plum Island Animal Disease Center, ARS, USDA, NY, USA; Oak Ridge Institute for Science and Education, PIADC Research Participation Program, Oak Ridge, TN, USA; Los Alamos National Laboratory, NM, USA; Department of Veterinary Population Medicine, University of Minnesota, Twin Cities, MN, USA; Department of Pathobiological Sciences, School of Veterinary Medicine, University of Wisconsin-Madison, Madison, WI, USA

## Abstract

Foot-and-mouth disease (FMD) is a highly contagious viral disease that severely impacts global food security and is one of the greatest constraints on international trade of animal products. Extensive viral population diversity and rapid, continuous mutation of circulating FMD viruses (FMDVs) pose significant obstacles to the control and ultimate eradication of this important transboundary pathogen. The current study investigated mechanisms contributing to within-host evolution of FMDV in a natural host species (cattle). Specifically, vaccinated and non-vaccinated cattle were infected with FMDV under controlled, experimental conditions and subsequently sampled for up to 35 days to monitor viral genomic changes as related to phases of disease and experimental cohorts. Consensus-level genomic changes across the entire FMDV coding region were characterized through three previously defined stages of infection: early, transitional, and persistent. The overall conclusion was that viral evolution occurred via a combination of two mechanisms: emergence of full-genomic minority haplotypes from within the inoculum super-swarm, and concurrent continuous point mutations. Phylogenetic analysis indicated that individuals were infected with multiple distinct haplogroups that were pre-existent within the ancestral inoculum used to infect all animals. Multiple shifts of dominant viral haplotype took place during the early and transitional phases of infection, whereas few shifts occurred during persistent infection. These insights into FMDV population dynamics have important implications for virus sampling methodology and molecular epidemiology.

## Introduction

Foot-and-mouth disease (FMD) is a highly contagious viral disease that affects wild and domestic even-toed ruminants [1, 2]. FMD is a major global concern for livestock owners and managers, and the disease has substantial impact on regulation of international trade in animal products [3]. The classical signs of disease include oral and pedal vesicles and erosions, often associated with lameness, pyrexia, and obtundation [2, 4]. The causative agent, FMD virus (FMDV) is extremely contagious and disseminates rapidly amongst susceptible animals. Although the disease is rarely fatal, FMD-endemic regions incur substantial economic burdens associated with production losses and disease control [5]. Sporadic outbreaks in countries that are normally free from FMD result in costly mediations including culling of large numbers of animals and animal movement restrictions, as well as massive economic losses due to implications on trade in animal products.

FMDV (family: *Picornaviridae*, genus: *Aphthovirus*) is a single-stranded RNA virus which exists in 7 defined serotypes; O, A, C, Asia-1, and Southern African Territories (SAT) 1-3. The FMDV genome encodes a total of 12 mature proteins translated from a single polyprotein coding region approximately 7 kilobases in length. The structural proteins VP1, VP2, VP3, and VP4 compose the capsid, with all but VP4 serving as major antigenic targets and taking part in receptor-mediated host cell entry [4, 6]. Positive selection has been identified predominantly within the capsid-encoding segments, leading to elevated amino acid replacement rates [7–9].

Existing population diversity and rapid, continuous mutation of circulating FMD viruses pose significant obstacles to its control and ultimate eradication. FMDV and other picornaviruses have some of the highest mutation rates measured, at approximately one mutation per genome per replication cycle (~7.8 × 10^-4^ nucleotides per copy) due to a low-fidelity RNA-dependent RNA polymerase [10–12]. Although estimates vary between studies and serotypes, FMDV nucleotide substitution rates are generally found to be higher within-host than between-host (transmission chains) [8, 13–15]. Furthermore, the within-host diversity of picornavirus populations has been shown to directly correlate with pathogenicity [16, 17]. The role of viral quasispecies swarms in the evolution of FMDV has been thoroughly described in tissue culture-based studies[18]; however, few studies have investigated this phenomenon in vivo in natural hosts[19–21].

In cattle infected with FMDV, the initial site of viral replication has been localized to focal regions of epithelium within the nasopharyngeal mucosa [22, 23]. The acute phase of disease lasts approximately one week, however, a substantial proportion of infected cattle become subclinical long-term carriers of the virus [24–26]. In carrier cattle, virus replication is restricted to epithelial cells of the nasopharynx [27–29]. The role of these animals in transmission of FMDV is controversial. However, the occurrence of FMDV carriers, particularly amongst vaccinated animals, has important ramifications concerning international trade and outbreak response measures [30, 31]. The FMDV carrier state has conventionally been defined by the presence of infectious virus in oropharyngeal fluids (OPF) >28 days past initial infection [32]. However, recent investigations have demonstrated that animals that clear infection can be differentiated from those that become carriers as early as 10 days post infection (dpi) for vaccinated and 21 dpi for non-vaccinated cattle [28, 33]. These findings have led to the definition of the transitional phase of FMDV infection, which corresponds to the temporal window during which FMDV is cleared from cattle that do not become carriers [28]. Transitional phase events thus comprise a critical stage of FMDV pathogenesis wherein pivotal virus-host interactions determine the outcome of the FMDV carrier state divergence [28, 34].

Detailed analysis of longitudinal virus samples obtained from individual hosts has increased understanding of antigenicity and important microevolutionary processes in RNA viruses such as hepatitis C virus and influenza virus [35–37]. A vivid example of the importance of events that take place during chronic infection is the case of *poliovirus* wherein the Sabin vaccine strain evolved pathogenicity in a patient over time [38, 39]. Recent publications examining the full-length FMDV genome have begun to elucidate the complexities of viral population dynamics and behavior through transmission events and within hosts [19, 21, 40, 41]. A detailed understanding of how FMDV changes over time both within hosts and through chains of transmission is key to understanding its pathogenesis and epidemiology.

The central aim of this study was to characterize changes that take place within the FMDV coding region across all stages of infection in vaccinated and non-vaccinated cattle. The duration and longitudinal nature of this study enabled novel characterizations of emergent FMDV variants as related to disease stage, vaccination status, and haplotypic linkage.

## Methods

### Experimental design

This investigation was based on samples collected from cattle included in a large-scale investigation of the FMDV carrier state, carried out at the Plum Island Animal Disease Center, New York. All procedures involving animals were carried out in accordance with the experimental protocol (protocol 209-15-R) that had been approved by the Plum Island Animal Disease Center Institutional Animal Care and Use Committee. Details of the animal experiments and sample collection have been described in previous publications [22, 28, 42]. In brief, both vaccinated (with a recombinant adenovirus-vectored FMDV serotype A vaccine [43]) and non-vaccinated cattle were infected with FMDV-A24 Cruzeiro through intra-nasopharyngeal inoculation [44] and monitored for up to 35 days post infection (dpi). All animals were subjected to daily clinical examinations, and analgesics and anti-inflammatory drugs (flunixinmeglumine, 1.1–2.2mg/kg; butorphanol tartrate, 0.1 mg/kg) were administered if needed to mitigate pain associated with severe foot-and-mouth disease. Animals were euthanized for tissue harvest at pre-determined time points throughout the study by intravenous injection of sodiumpentobarbital (86 mg/kg).

Three distinct periods were used to define the progression of infection in individual animals: early, transitional, and persistent periods, which have different temporal boundaries in non-vaccinated and vaccinated animals. In non-vaccinated cattle, the early phase (1-9 dpi) corresponds to the clinical phase of disease with viremia and systemic generalization of infection. These animals recover from clinical disease and either clear infection completely during the subsequent transitional phase (10-21 dpi) or maintain a subclinical infection of the nasopharynx through the persistent phase (>21 dpi). All vaccinated cattle included in this investigation were protected against clinical FMD and systemic infection. For vaccinated cattle, the early phase of infection comprised primary infection of the nasopharyngeal mucosa, and associated shedding of low quantities of virus in oral and nasal secretions. Similar to non-vaccinated animals, a subset of the vaccinated cattle cleared infection during the transitional phase, which in this cohort was defined as 7-14 dpi, based on distinct infection dynamics compared to non-vaccinated animals [28]. For both vaccinated and non-vaccinated cattle, the persistent phase of infection (>15 dpi and >21 dpi, respectively) consisted of subclinical persistence of infectious FMDV in the nasopharyngeal mucosa.

### Virus inoculum

The virus inoculum used to infect cattle in the current investigation was generated from a field-derived strain of FMDV A24 Cruzeiro, which was grown in BHK-21 cells (ATCC, Manassas, VA) and subsequently passaged twice in cattle. In passage 1, two cattle were inoculated through intra-epithelial injection in the tongue. At 48 hours post inoculation, vesicular lesions harvested from the tongue and feet of the infected cattle were used to generate a pooled virus suspension. This suspension was used to infect a second cohort of three cattle through tongue inoculation. At 48 hours post inoculation, a second pooled virus suspension was generated from tongue and foot vesicles of this second cohort. This second passage suspension was aliquoted and stored at −70°C. All cattle included in the current investigation were inoculated with 10^5^ BTID_50_ (50% infectious doses titrated in bovine tongue epithelium) of the second passage virus.

### Samples

Samples collected primarily to monitor disease progression included oral and nasal swabs, serum and OPF (sampled using a probang cup [45]). Additionally, tissue distribution of virus was investigated in samples obtained at necropsy examinations which were performed at predetermined experimental endpoints, regardless of disease progression. All samples were screened for the presence of FMDV genomic RNA and infectious virus. The detailed approach and outcome of these investigations have been previously published [22, 28, 42].

For the current investigation, fifty-two specimens from 13 cattle were selected for FMDV sequencing and analysis (S1 Fig). These samples were selected based upon positive isolation of FMDV and included samples from early, transitional, and persistent phases of infection from both vaccinated and non-vaccinated cattle. This sample set included 41 samples from live animals and 11 postmortem tissue samples. Tissue samples collected postmortem were either vesicular lesions (Ves) or nasopharyngeal mucosa (Np).

### Virus sample processing and sequencing

Each sample was singly passaged in LFBK-αvβ6 cells in order to ensure sufficient virus for subsequent steps. Thirty-two samples were sequenced using a sequence-independent single primer amplification method using the Illumina MiSeq platform as previously described [46]. An additional twenty samples as well as the inoculum were sequenced on the Illumina NextSeq 500. For the NextSeq-sequenced samples, genomic FMDV RNA was extracted using the MagMAX RNA Isolation Kit (Thermo Fisher Scientific) on the KingFisher particle processor. Three overlapping amplicons were reverse-transcribed (SuperScript III, Thermo Fisher Scientific) and synthesized from genomic RNA using primer pairs TGGTGACAGGCTAAGGATG / GCCCRGGGTTGGACTC (5’-UTR – 2A), AGTGTACAACGGGACGAGTAAGTAT / TTGCTCTCTCAATGTACTCACTCAC (VP1 – 3A), and TGGCAATGTTTCAGTACGACT / CGCGCCTCAGAAACAGT (2C – 3’-UTR). Amplicons were quantified using the Qubit 2.0 fluorometer / RNA BR/HS Assay Kit (Thermo Fisher Scientific) and normalized accordingly. Libraries were constructed with the Nextera XT DNA Library Prep Kit (FC-131-1096, Illumina, San Diego, California, USA). All reads were finished using CLC Genomics Workbench v. 9.5 (www.qiagenbioinformatics.com), including primer removal and quality-trimming. All reads were mapped to the FMDV-A24 Cruzeiro reference genome. The sequence-independent amplification method (32 samples) provided relatively low coverage in regions, thus a minimum coverage was set at 10 reads for base calling across all samples, including the targeted amplicon sequenced samples. For this reason, virus consensus sequences only (*no* subconsensus data) were analyzed from the experimental animal samples. The inoculum was the only material for which deep sequencing data was utilized.

### Sequence analysis: alignment, statistics, variation

Alignments and pairwise distances were determined in MEGA 7.0 [47] and Geneious 7.1 (www.geneious.com [48]). Substitutions (nucleotide differences) were tabulated for each sampled virus consensus sequence as compared to that of the inoculum consensus. Step-wise rates were also calculated by counting substitutions between successive samples divided by the time elapsed between samples. Step-wise rates were averaged across groupings according to vaccination status and phase of infection. For instances in which there were multiple samples from the same animal on the same date (differing only by sample type), total substitutions were averaged solely for rate comparisons. The CLC Genomics Workbench low frequency variant tool was used to determine site variation present in the inoculum at or above 2% with 0.75 strand-bias filter from 10.4 million reads mapped to its own consensus (200k average and 73k minimum coverage across the protein coding region).

### Phylogenetic analysis

The phylogenetic relationship between consensus viruses was modeled using PhyML 2.2 software (www.atgc-montpellier.fr/phyml, [49]). An inoculum-rooted maximum likelihood tree was constructed from 2000 sampled trees with the parameters: consensus threshold 0.5; model HKY98, 4-bin gamma; Ts/Tv 4.

### FMDV capsid protein modeling

Homology modeling of the inoculum was performed using PDB: 1FOD template with Prime Homology modeling module of Schrödinger Maestro v.11[50]. Residues were mutated using Maestro workspace and structures were minimized with Prime using the VSGB solvation model. Protein structures were rendered using Schrödinger Maestro, PyMOL and APBS plugin for surface electrostatics. Antigenic sites (A-1 to A-5) were defined as described in Fry et al., 2005 [51].

### Data availability

The 52 FMDV sample sequences are available in GenBank, accession numbers MH426523-74. Inoculum (SRA) reads are accessible in GenBank at SRP149342.

## Results

### Clinical studies

This study was based upon analyses of virus samples harvested from animals that were part of a large-scale experimental investigation of the FMDV carrier state divergence in cattle. The clinical outcomes of the experiments, including monitoring of infection in live animals and determination of the tissue distribution of virus through defined phases of infection, have been previously published [22, 28]. In brief, vaccinated and non-vaccinated cattle were infected with FMDV strain A24 Cruzeiro through simulated-natural inoculation and were monitored through 35 days. All vaccinated cattle were protected from clinical FMD, whereas all non-vaccinated cattle developed generalized FMD within 3-5 dpi. Vaccinated cattle were all subclinically infected, and the occurrence and characteristics of persistent FMDV infection were similar between vaccinated and non-vaccinated cohorts.

### Variation present in the inoculum

In order to approximate the diversity of FMDV circulating in an outbreak, the A24 Cruzeiro virus inoculum used to experimentally infect all animals in these experiments was composed of pooled virus acquired from vesicles from three cattle inoculated with the same ancestral strain (described in methods). The heterogeneity of the inoculum was investigated with ultra-deep sequencing of the full genomic coding region. Across 10.4 million mapped reads, the variant analysis identified 33 minority nucleotide variants present between 2.1 and 48.9%, distributed at unique sites across the protein coding region in all but the smallest genes, 2A and 3B (Fig 1). Limitations of sequencing methodologies (e.g. inconsistencies in coverage) dictated that consensus-level investigation of virus samples would be the most appropriate level at which to analyze genetic data. In total, 20 (60.6%) of inoculum subconsensus variants subsequently emerged as the majority (consensus) in viral isolates from animal samples within this study. Based upon phylogenetic relationships of the consensus sequences from animals’ samples and synchronous nucleotide changes (detailed below), distinct clades were inferred within the inoculum. The clade-characteristic single nucleotide polymorphisms (SNPs) present within the subconsensus variation of this complex inoculum were considered to have greater diversity than could be attributed to a conventional quasispecies swarm. On this basis, the heterogeneous inoculum is described herein as a super-swarm in order to illustrate the constituent complexity derived from several individual swarms (Fig 1).

**Fig 1.**
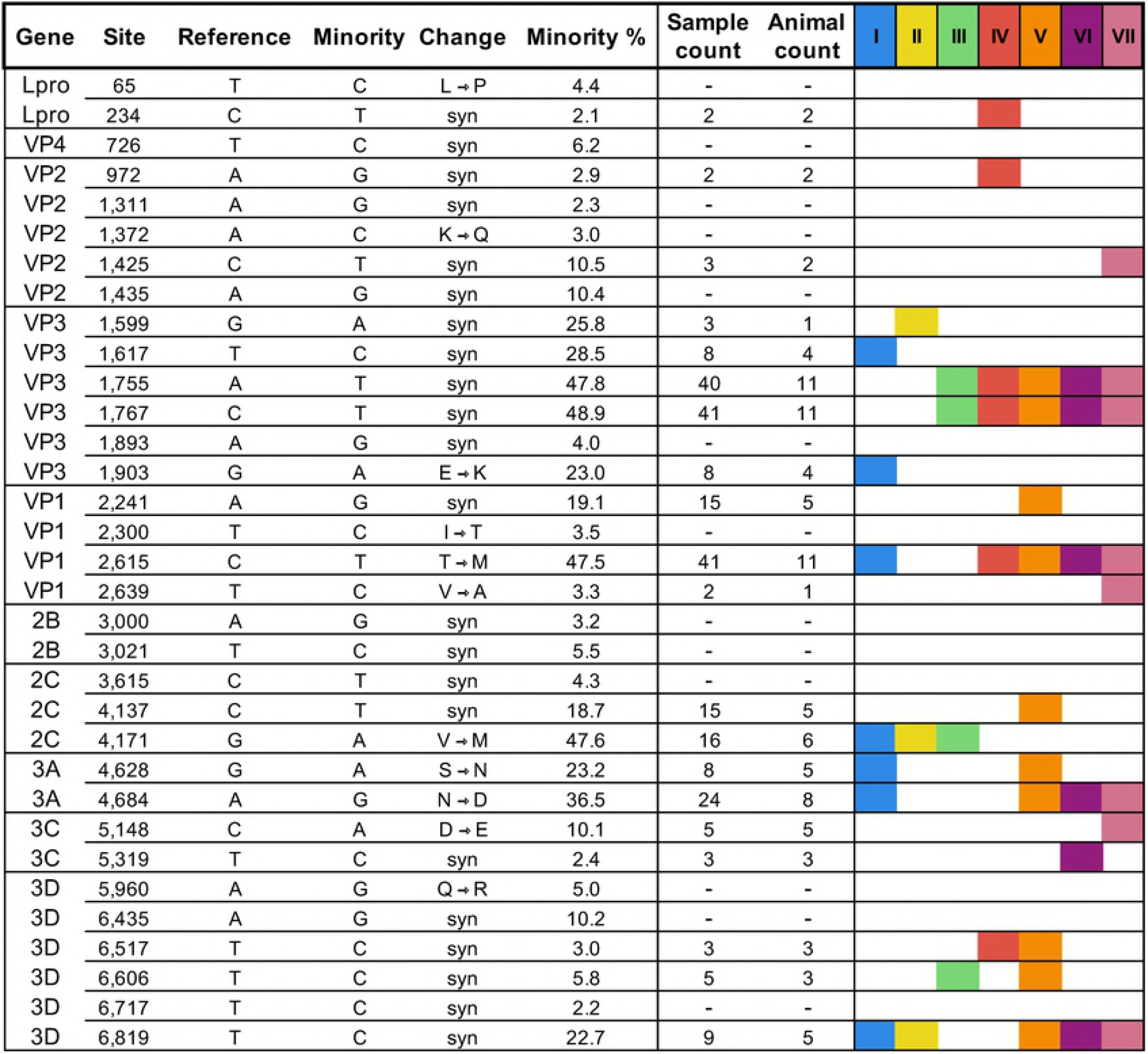
Subconsensus variants present above 2% frequency in inoculum ultra-deep sequencing. The reference used was the inoculum consensus. Sample count indicates the number of sample consensus sequences (of 52 total) that encoded each nucleotide found at the minority (2 - 49%) level in the inoculum deep sequence. Animal count indicates the number of individual hosts that provided sequences with these variants of the 13 total. If no value is indicated (-), no sample sequence encoded the minority nucleotide at the consensus level. The presence of each specific site change in at least one clade member (see phylogeny, Fig 4) are indicated by associated colors.

### Gene-specific comparisons of nucleotide substitutions

Consensus-level nucleotide changes were characterized in 52 virus samples obtained from 13 experimentally infected cattle (7 non-vaccinated and 6 vaccinated), through the experimental period which lasted up to 35 days. A subset of animals was euthanized for harvest of tissue samples at pre-determined time points during early and transitional phases of infection and therefore did not contribute to the investigation of persistent phase viruses. Samples included in the analysis were nasal fluid, saliva, oropharyngeal fluid (OPF), serum, vesicle epithelium, and nasopharyngeal tissue samples. Comparing the 52 consensus-level virus sequences to the parental inoculum consensus sequence revealed 545 total nucleotide changes distributed across 168 sites, with at least one substitution in each viral protein’s coding sequence (Table 1). The coding sequences for capsid proteins VP3 and VP1 had the highest proportions of substituted sites at 3.8 and 3.5%, respectively (Table 1). Additionally, VP3 and VP1 contained the highest numbers of nonsynonymous changes and each coding segment contained two codons with nonsynonymous changes at different codon positions. Multiple amino acid substitutions were found in coding segments of capsid genes of all serially sampled animals. The lowest proportion of nucleotide change occurred in VP4 (0.4%) and 3D (1.6%).

**Table 1.**
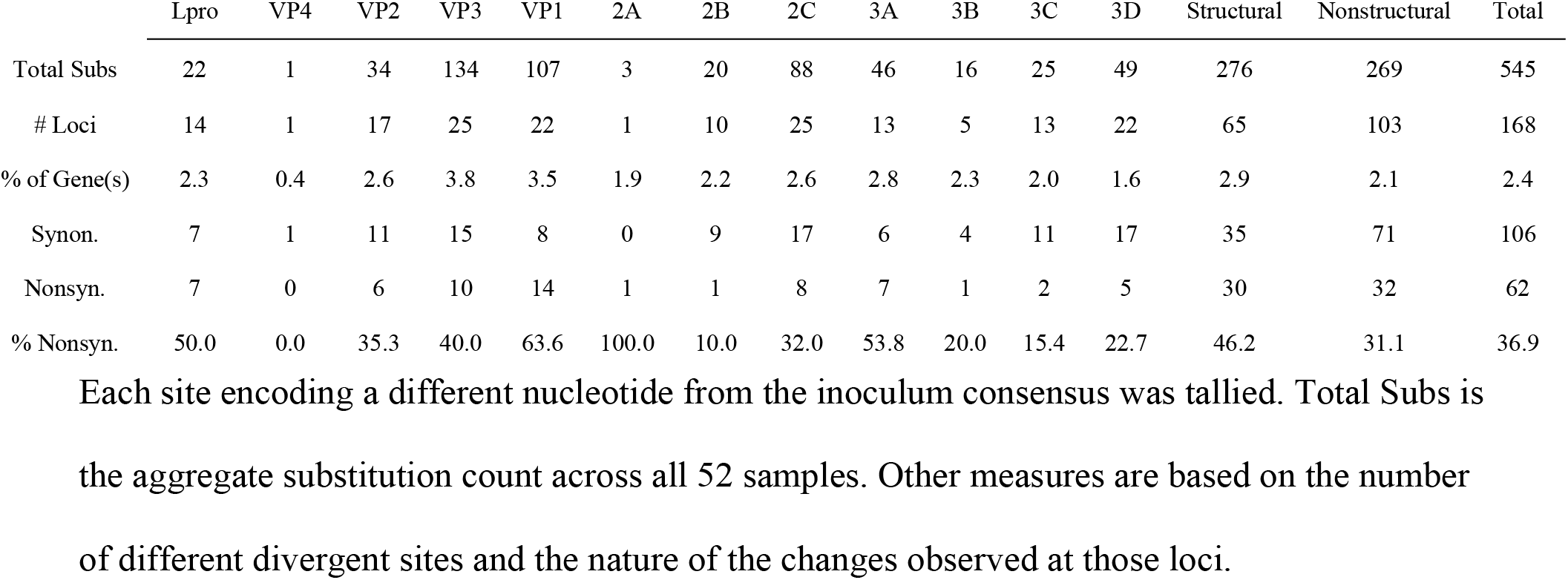
Distribution of consensus-level nucleotide substitutions in sample viruses across the FMDV coding region in 52 sampled viruses.

|  | Lpro | VP4 | VP2 | VP3 | VP1 | 2A | 2B | 2C | 3A | 3B | 3C | 3D | Structural | Nonstructural | Total |
| --- | --- | --- | --- | --- | --- | --- | --- | --- | --- | --- | --- | --- | --- | --- | --- |
| Total Subs | 22 | 1 | 34 | 134 | 107 | 3 | 20 | 88 | 46 | 16 | 25 | 49 | 276 | 269 | 545 |
| # Loci | 14 | 1 | 17 | 25 | 22 | 1 | 10 | 25 | 13 | 5 | 13 | 22 | 65 | 103 | 168 |
| % of Gene(s) | 2.3 | 0.4 | 2.6 | 3.8 | 3.5 | 1.9 | 2.2 | 2.6 | 2.8 | 2.3 | 2.0 | 1.6 | 2.9 | 2.1 | 2.4 |
| Synon. | 7 | 1 | 11 | 15 | 8 | 0 | 9 | 17 | 6 | 4 | 11 | 17 | 35 | 71 | 106 |
| Nonsyn. | 7 | 0 | 6 | 10 | 14 | 1 | 1 | 8 | 7 | 1 | 2 | 5 | 30 | 32 | 62 |
| % Nonsyn. | 50.0 | 0.0 | 35.3 | 40.0 | 63.6 | 100.0 | 10.0 | 32.0 | 53.8 | 20.0 | 15.4 | 22.7 | 46.2 | 31.1 | 36.9 |

Each site encoding a different nucleotide from the inoculum consensus was tallied. Total Subs is the aggregate substitution count across all 52 samples. Other measures are based on the number of different divergent sites and the nature of the changes observed at those loci.

### Sequence alignment

The consensus virus sequences obtained within the first five days of infection (acute phase) were highly similar to the inoculum consensus. Substitutions in these early phase samples were largely shared across multiple animals and all of those shared substitutions were present at frequencies greater than 5% in the inoculum (Fig 1). For example, inoculum subconsensus variants T1617C and G1903A were each identified at consensus level in the earliest samples in 4 different animals, whereas variants A1755T, C1767T, and C2615T were each present in the earliest consensus viruses from 11 of the 13 total hosts (S1 Fig). At later stages of disease, most new consensus-level changes present in samples were not among those detected at subconsensus level in the inoculum (>2%) (Fig 1 and S1 Fig). The general trend across sample consensuses was an increased divergence from the inoculum over time; however, this did not occur consistently in all animals (Fig 2a and S1 Fig).

**Fig 2.**
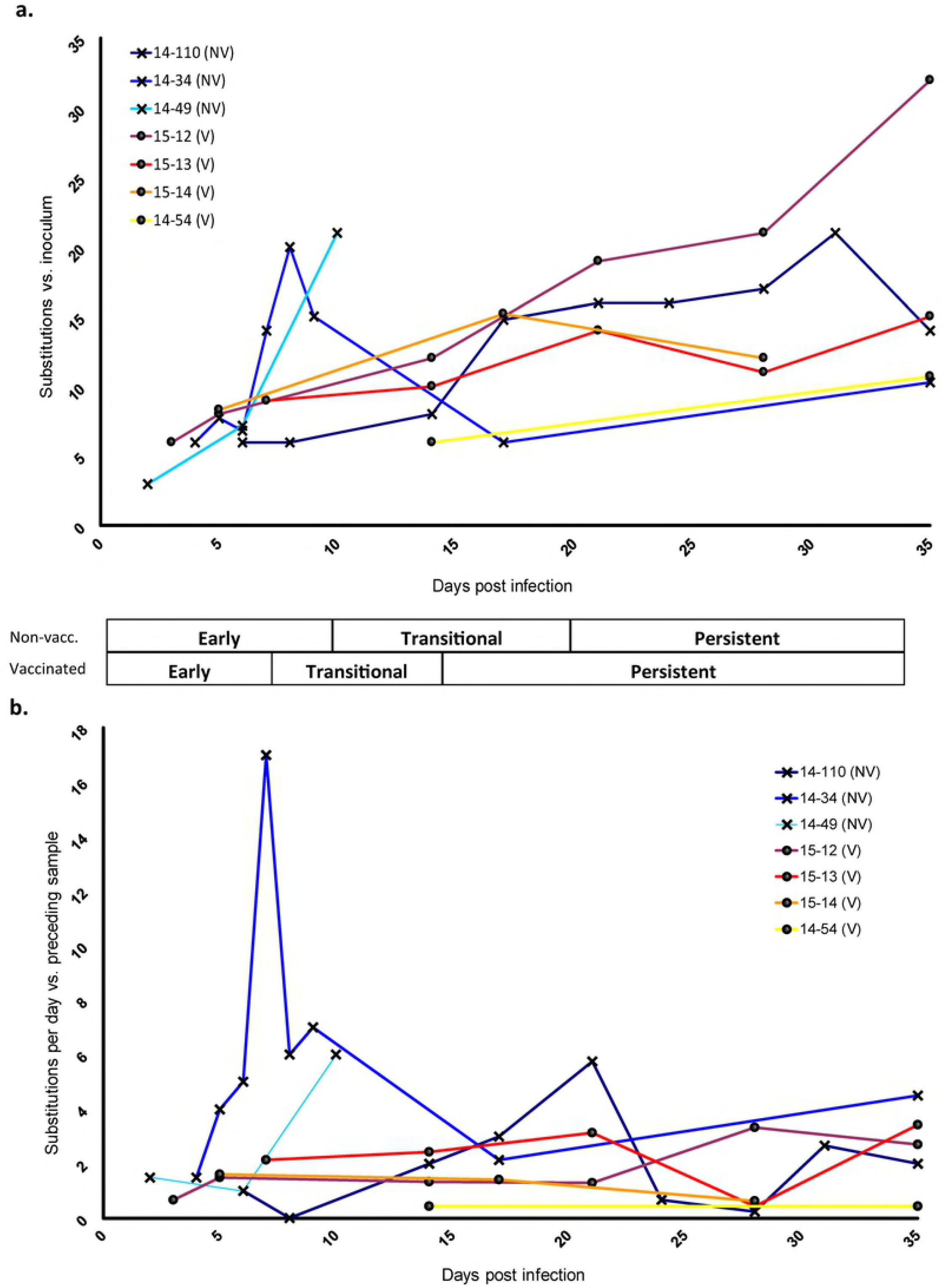
FMD virus change over the course of infection. a) The total number of nucleotide substitutions present at the consensus level at each sampled time point compared to inoculum consensus b) Step-wise changes per day. The number of substitutions (nucleotide differences between each sample consensus and the previous sample consensus) divided by the number of elapsed days between samples plotted against each sample time since initial infection. Values from concurrent samples from the same animal (different tissues) were averaged. Overlapping data points were slightly offset for clarity. Early, Transitional, and Persistent phases of infection are described in methods.

### Nucleotide substitutions over time

Quantitation of consensus-level nucleotide changes within each animal during infection as compared with the inoculum consensus indicated a variable, but overall positive slope of sequence change over time (Fig 2a). However, reversions (down-turning curves) were common and occurred at least once in four of the seven serially-sampled animals. In order to account for potential bottleneck events and adjust for elapsed time between samplings, nucleotide changes from sample to sample over elapsed time were also plotted (Fig 2b). While the sequences almost always differed at multiple sites between sampling dates, the rates of change were relatively stable in vaccinated hosts over time as compared to non-vaccinated. Averaged substitution rates from sequential samples within the distinct phases of disease demonstrated that non-vaccinated animals had higher rates of sequence change compared to vaccinated animals during early (0.24 vs. 0.08 substitutions/site/year) and transitional (0.17 vs. 0.09 subs/site/yr) phases (Table 2), though these differences were not statistically significant. During the persistent phase, the substitution rates were similar for non-vaccinated and vaccinated cohorts (0.11 versus 0.10 subs/site/yr). A gradual accrual of substitutions, consistent with canonical evolutionary processes, was commonly observed as demonstrated by animal 14-34 between 17dpi-35dpi and animal 14-110 from 21dpi-35dpi (Fig 2a). However, the unprecedented rates as measured alongside substitution patterns between consecutive samples consisting of synchronous changes across the genome suggested a process that was more complex than independent nucleotide substitution. Specifically, sample-to-sample substitution patterns that represented irregular jumps in substitution rates (Fig 2) were highly suggestive of linkage between variant nucleotides representing distinct genotypes. The rapidity and scope of this phenomenon were exemplified in animal 14-34, where 17 nucleotide changes in 7 genes, spanning Lpro to 3D, differed at the consensus level within a 24-hour period (6 to 7 dpi; Fig 2 and S1 Fig). Subsequently, a set of 17 (mostly revertant) nucleotide changes differed between 9 and 17 dpi in this same animal, suggesting shifts in dominance between variant viral genomes.

**Table 2.** Nucleotide substitution rates averaged across non-vaccinated and vaccinated cattle.

|  | **Early** | **Transitional** | **Persistent** |
| --- | --- | --- | --- |
| **Average Substitution Rate - Non-vaccinated Cattle** |  |  |  |
| Substitutions /day | 4.55 | 3.22 | 2.02 |
| Substitutions /site /day | $6.49 \times 10^{-4}$ | $4.60 \times 10^{-4}$ | $2.88 \times 10^{-4}$ |
| Substitutions /site /year | $2.37 \times 10^{-1}$ | $1.68 \times 10^{-1}$ | $1.05 \times 10^{-1}$ |
| **Average Substitution Rate - Vaccinated Cattle** |  |  |  |
| Substitutions /day | 1.48 | 1.73 | 1.93 |
| Substitutions /site /day | $2.11 \times 10^{-4}$ | $2.47 \times 10^{-4}$ | $2.75 \times 10^{-4}$ |
| Substitutions /site /year | $7.71 \times 10^{-2}$ | $9.01 \times 10^{-2}$ | $1.00 \times 10^{-1}$ |

### Phylogenetic associations

Phylogenetic analysis led to delineation of seven distinct clades (I-VII, Fig 3). A unique set of variant nucleotides defined each clade with a minimum of 2 substitutions differentiating clades IV, VI, or VII (red, purple, and pink) and a maximum of 8 nucleotide changes differentiating clade III members (green) from all other viruses. Importantly, the clade-characterizing nucleotide changes identified in the animal samples were also present as subconsensus variants within the inoculum, wherein they averaged 27.2% frequency (2.4 – 48.9%) (Fig 1). As expected, when using the inoculum as the root, the earliest sampled virus sequences were basal, while later samples tended to lay furthest from the root. Yet, there were numerous outliers such as the 10 dpi nasopharyngeal tissue sample from animal 14-49 and the 8 dpi saliva sample from animal 14-34 (both in clade III), for which the long branch lengths indicated that substantial changes occurred at these intermediate time points. Viruses were initially expected to cluster according to individual animal IDs, with sequential progression of divergence and later samples descending from earlier in-host viruses. However, that pattern was not consistent. Instead, virus samples were often found to share a closer phylogenetic relationship with those obtained from other animals (Fig 3). For example, clades I and V include viruses derived from 4 and 5 different hosts, respectively.

**Figure 3.**
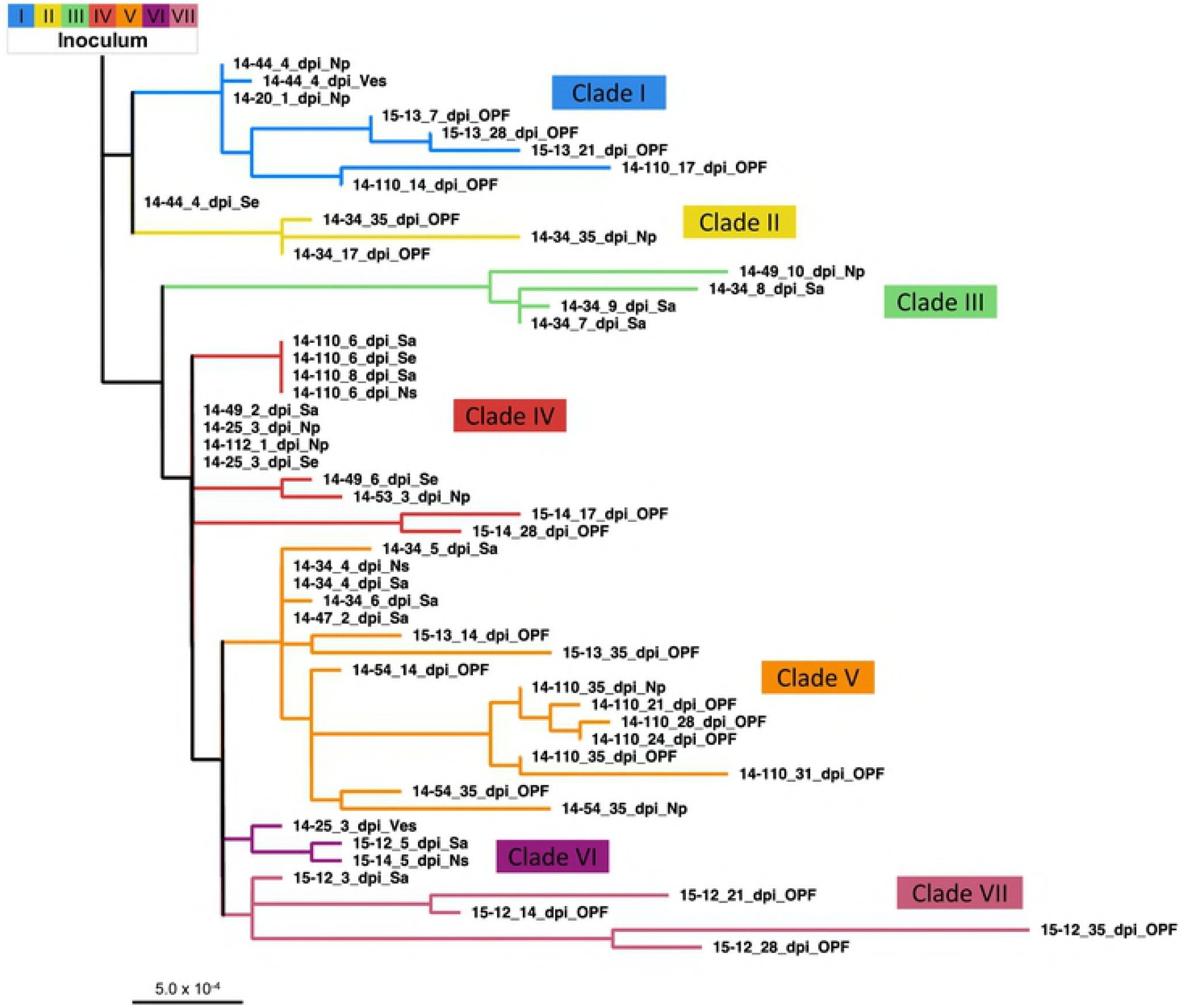
Phylogenetic relationships between sampled virus consensuses. An inoculum-rooted maximum likelihood tree was created using PhyML. Branch lengths are proportional to nucleotide differences. Clades have been delineated based upon the clusters present here. Abbreviations: dpi – days post-infection, Na – nasal secretion, Np – nasopharyngeal tissue, OPF – oropharyngeal fluid, Sa – saliva, Se – serum, Ves – epithelial vesicle

### Clade shifts within animals

A clade shift was defined by the consensus sequence obtained at a given time point sharing a closer phylogenetic relationship to a different group of viruses as compared to the preceding sample from the same animal. Clade shifts took place in all serially-sampled cattle except animal 15-54, from which only three samples from the transitional and persistent phase were available for analysis. There were also instances of viruses of different clades coexisting in distinct samples collected at the same time point from an individual host (e.g. 14-25 at 3 dpi; Fig 4). However, there was no consistent pattern of association between the sample type and the clade of virus identified (Fig 4). Interestingly, clade reversion (return of a clade previously dominant within the same host) only occurred in vaccinated animals (animals 15-12 and 15-13); thus, in non-vaccinated animals, once a clade was cleared to below consensus, it did not return within the course of the study. There was no clear trend for any specific clade to dominate early infection. Rather, viruses belonging to five of the seven clades were identified in samples collected from the earliest samples (Fig 4).

There were several unique attributes to the clade shifts that occurred during the transitional phase of infection. Overall, there was a relatively high quantity of clade shifts during this period; all 7 clades and every animal sampled during this period manifested at least one clade shift. Clades III and VI were only detected during a relatively small temporal window spanning the end of the early phase through the transitional phase of infection; both of these clades were extinguished during the transitional phase. There was a strong tendency for consensus virus sequences not to undergo clade shifts during the persistent phase; however, individual consensus nucleotide changes continued throughout this period and were substantial in some animals (Figs 2 and 4). Only two clade shifts were identified during the persistent phase of infection, and both occurred in (vaccinated) animal 15-13 which had a shift from clade V to I and then back to V (Fig 4). Although 5 of the 7 clades were identified during persistent infection, clade V was ultimately overrepresented in the final 35 dpi samples, present at the consensus level in 3 of the 5 cattle sampled at this late time point.

**Fig 4.**
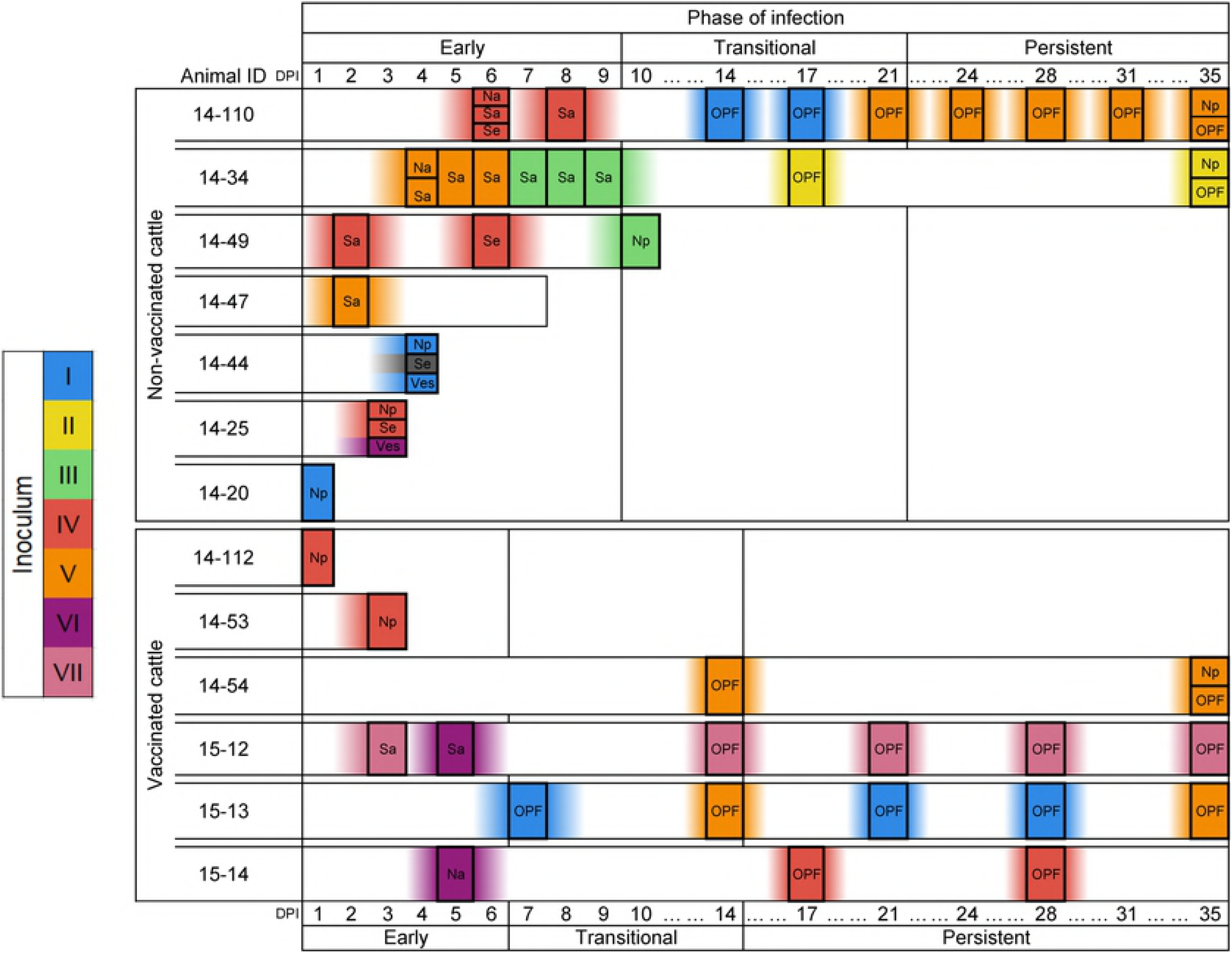
Consensus virus identity across infection. Boxes along each animal’s timeline indicates a virus sample. Colors and associated Roman numerals indicate clades based on inferred phylogenetic relationships (see Fig 4). Inoculum at left has been hypothesized as the source of the seven variant viruses detected. Abbreviations: DPI – days post-infection, Na – nasal secretion, Np – nasopharyngeal tissue (necropsy), OPF – oropharyngeal fluid (probang cup sample), Sa – saliva, Se – serum, Ves – epithelial vesicle. Three phases of FMDV infection were used to define progression of infection in individual animals: early, transitional, and persistent periods; temporal boundaries for the phases differ in vaccinated and non-vaccinated animals, as described in methods.

### Structural analysis

The FMDV capsid surface proteins VP3, VP2 and VP1 contain several well defined antigenic sites that are targeted by the antibody-mediated host response. Modeling and structural analysis of the virus capsid was performed in order to correlate viral genomic changes with associated changes in virus capsid structure. This investigation focused on the detection of amino acid substitutions present at sites, previously determined to be antigenically significant [51]. Structural modeling of the FMDV-A_24_ Cruzeiro capsid protomer including proteins VP1, VP2, and VP3 indicated that major antigenic sites and adjacent regions harbored a substantial proportion of variable residues (Table 3, Fig 5). As VP4 lines the capsid interior, it was excluded from this part of the analysis. The two most common amino acid substitutions that became fixed in the animal samples were in VP1 sites 144 and 147 (Table 3). Notably, these amino acid residues are within the receptor-binding site in the antigenically dominant G-H loop of VP1.[52, 53] (Fig 5). A threonine to methionine substitution at VP1 site 147 was the most frequently observed amino acid change (Table 3). This substitution preserves the local hydrophobic pocket that is important for host receptor binding and antibody neutralization (Fig 5b). Deep sequencing of the inoculum virus indicated that methionine was present at this position at a frequency of 47.5% prior to exposure to animals, which may account for this commonly observed substitution. In contrast to this finding at VP1 147, arginine at VP1 144 was present at <2% frequency in the inoculum, yet a serine to arginine substitution at this position was found in 20 out of the 52 sampled viruses, all obtained ≥7 dpi (Table 3). Other substitutions that occurred at sites adjacent to antigenically relevant locations and were found in samples from multiple animals were threonine to lysine at VP3 175 (Fig 5a, c) and glutamic acid to lysine or glycine at VP3 131 (Fig 5a). The VP3 T175K substitution was present only in clade VI viruses and was found in samples from animals 15-12 and 15-14 during early infection. Interestingly, the VP1 S144R and VP3 T175K substitutions did not co-exist in any haplotype (Table 3), suggesting that the lysine at VP3 175 may represent an accommodation to the atypical serine at VP1 144 that was present in the inoculum and the majority of early phase animal samples. VP1 199 is located at the VP1 C-terminus (Fig 5a, d) and has also been identified as critical for integrin binding [54]. An aspartic acid to glycine substitution at this position was found in two samples obtained at 10 and 14 dpi, respectively. This D199G substitution causes a loss of local hydrogen bonding (Fig 5d) and can affect the conformation and surface charge of the antigenically critical VP1 C-terminus (Fig 5e, middle panel). Other substitutions with potential influence on the capsid structure were VP2 residues 82, 88, and 131. Substitutions that occurred at these sites were transient and disappeared during later stages of infection, highlighting the dynamic nature of virus evolution and host pressure.

**Table 3.**
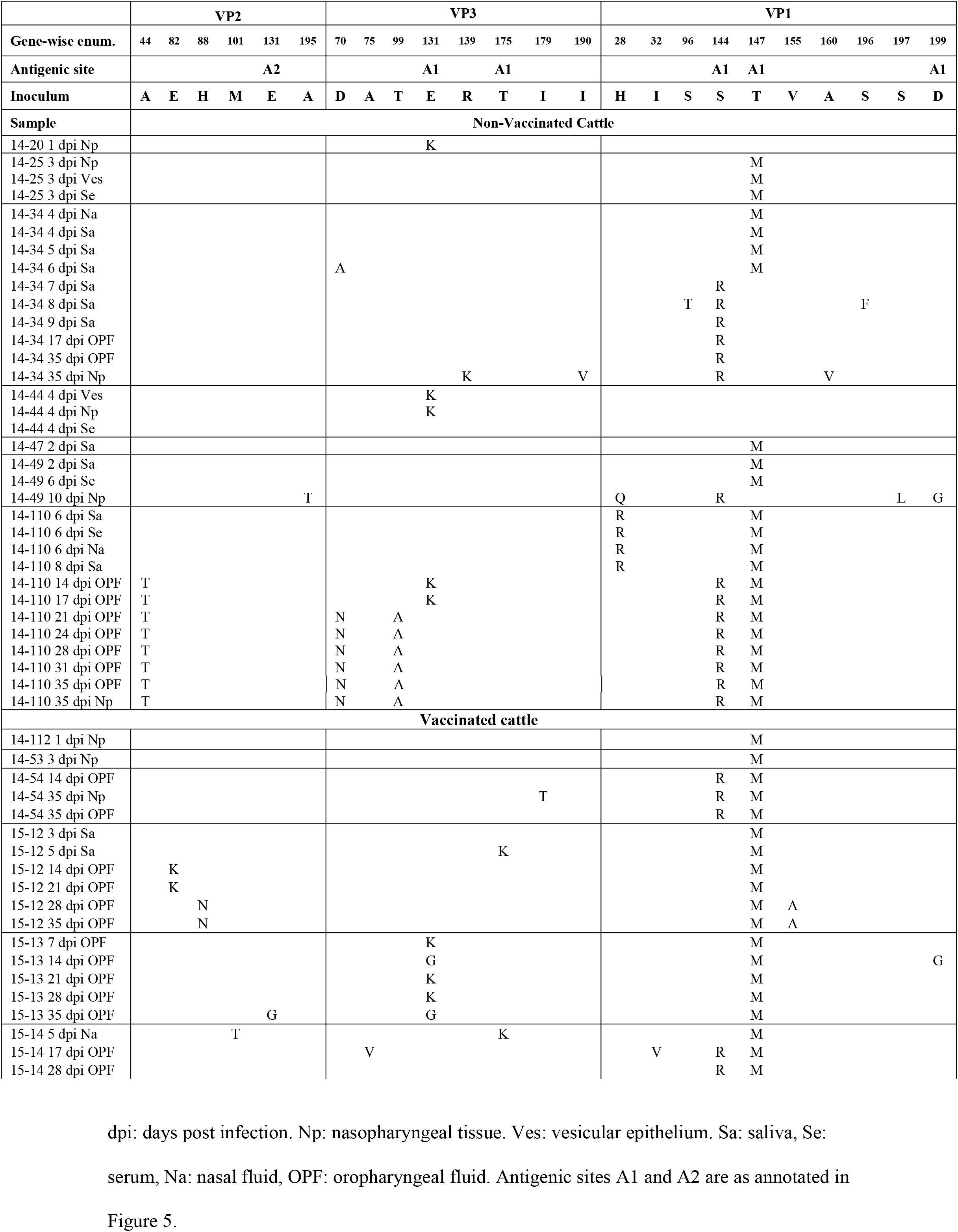
Amino acid substitutions in FMDV capsid surface proteins

**Fig 5.**
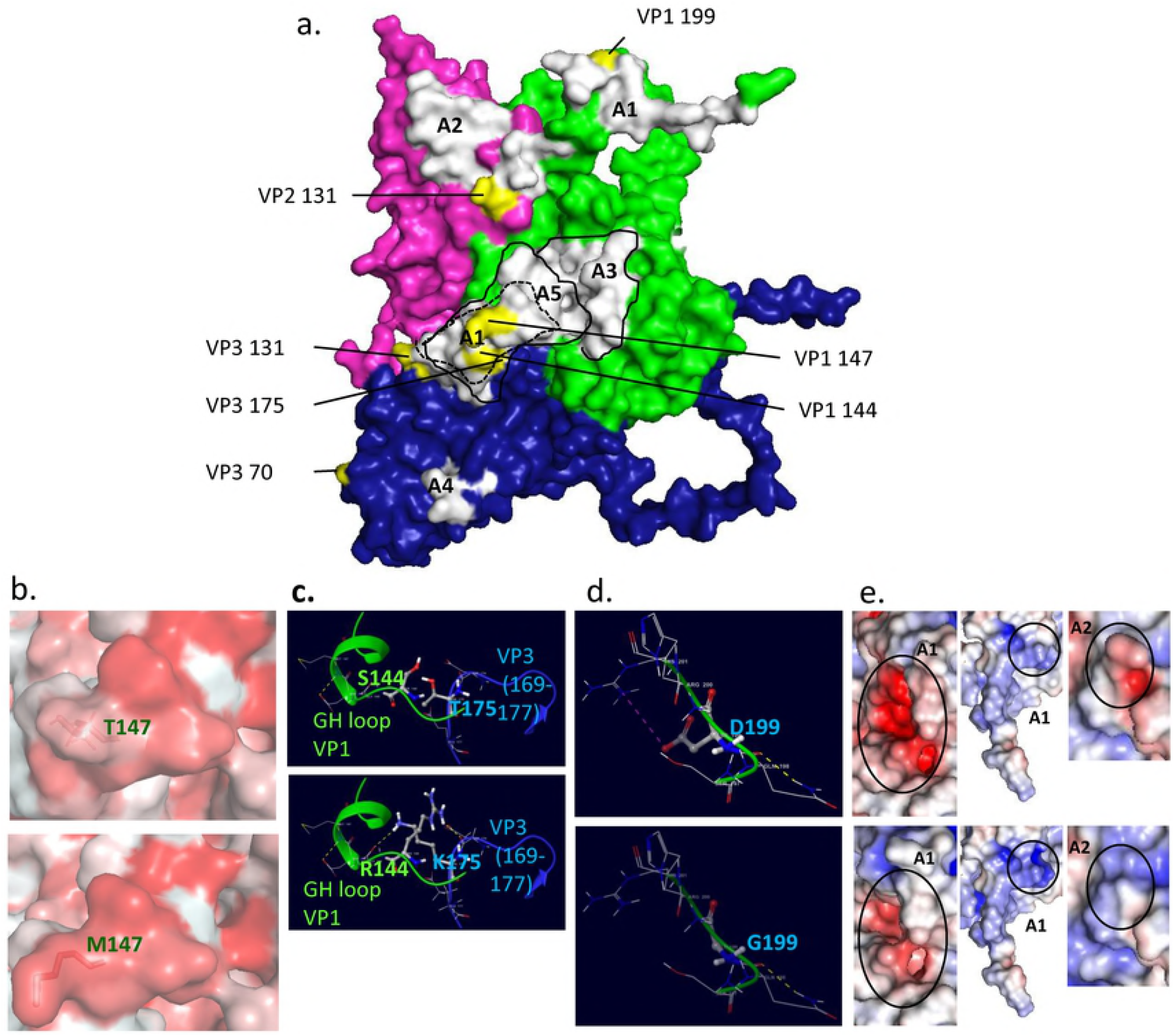
Molecular modeling of virus capsid. a) Surface representation of exposed (outer) surface inoculum virus capsid protomer. VP1, VP2 and VP3 are green, magenta and blue with major antigenic sites marked as A1 to A5. Substitution sites are marked in yellow and labeled with gene-wise enumeration. b) Hydrophobic pocket surrounding residue (red intensity correlates with stronger hydrophobicity) VP1 147 (shown as sticks). Inoculum virus (upper panel) and the majority of sampled viruses (lower panel) exhibit conserved hydrophobicity. c) Polar bonding network involving GH loop residue VP1 144 and residue VP3 175 in inoculum virus (upper panel) and viruses with a threonine to lysine substitution at 175 or other viruses with an arginine substitution at 144 (lower panel). Secondary structure and residue labeling color code indicates corresponding capsid protein, blue for VP3 and green for VP1. d). Polar bonding involving residue 199 of VP1 C-terminus in inoculum virus (upper panel) and virus sample with a glycine substitution at this site (lower panel), green: VP1, blue: VP3. e). Electrostatic surface of major antigenic sites A1 and A2 in inoculum (upper panels) and sampled viruses (lower panels), blue, white and red colors indicate positively charged, neutral and negatively charged residues, respectively. Left panels represent surface change at A1, GH loop region and middle panels represent surface change at A1, C-terminus region (VP1 199). Right panels show surface charge gain adjacent to A2 (VP2 131).

## Discussion

In order to investigate FMDV evolution within natural hosts, cattle were experimentally infected with a complex, heterogeneous inoculum by a simulated natural route and progeny virus samples were collected over the course of 35 days post inoculation. Analysis of the virus consensus sequences obtained during different phases of infection suggested that at least two distinct processes contribute to within-host FMDV genomic changes: 1) conventional molecular evolution characterized by selection acting upon individual, *de novo* nucleotide substitutions and/or resultant amino acid changes, and 2) emergence and regression of full-genome minority haplogroup members over the course of infection.

The current study provides a detailed demonstration of the phenomenon of the multiplicity of founding infections and emergence of subconsensus variants as a common mechanism of FMDV evolution in natural hosts through all stages of infection. These data also demonstrate that consensus-level nucleotide changes detected during early infection were predominantly present at subconsensus level in the inoculum. The frequent haplotype shift observed within hosts is consistent with previous investigations of FMDV molecular evolution that have demonstrated that multiple divergent genotypes can be maintained within a host for extended periods of time. Specifically, strong evidence has been presented of distinct FMDV subpopulation co-existence in water buffalo (*Bubalus bubalis*) [13, 55] and cattle [56] during persistent infection. Similarly, African cape buffalo (*Syncerus caffer*) can simultaneously carry up to three FMDV serotypes for as long as 185 days after initial infection [57]. Additionally, a separate experimental study demonstrated that FMDV diversity was regularly maintained through transmission chains with specific variants reaching consensus in different acute-phase samples [19].

In the current study, the diverse virus subgroups present in the inoculum established multiple founder infections and individual haplotypes subsequently emerged at the consensus level at different times in different hosts. Phylogenetic analysis placed viruses from different animals into shared clades, supporting the hypothesis that these viruses (clade members) shared common ancestry present in the inoculum. If this phylogeny had occurred purely by drift and selection of non-linked nucleotide differences, the boundaries of each clade would be expected to be largely defined by substitutions unique to each animal. In contrast, 5 of the 7 clades contained viruses sampled from multiple animals.

Comparing virus isolates collected at the same time and from the same animal but from different anatomic locations provided examples of 3 distinct phenomena of within-host diversity: 1) identical consensus sequences across samples, 2) within-clade divergent sequences, and 3) viruses belonging to different clades. This range of possibilities demonstrates that at any specific time during infection, the virus population was evolving at both the individual nucleotide and haplotypic levels and that this evolution occurred independently at different sites within the same animal. The observed phenomenon of within-host diversity may have implications for various aspects of pathogenesis and transmission. In particular, chains of transmission may be affected, since only viruses at certain locations are likely to be directly transmitted to other animals (e.g. nasal swabs and lesions, but not serum or OPF).

This investigation included samples from both vaccinated and non-vaccinated cattle. As previously reported [28], there were striking differences in infection dynamics between the two cohorts; while all non-vaccinated animals had a phase of fulminant clinical disease with systemic generalization, infection in vaccinated cattle was subclinical and restricted to the upper respiratory tract. These differences in infection dynamics correlated with contrasting evolutionary processes taking place in the different cohorts of cattle. While the substitution rates measured in the present study were not interpreted as representing novel or fixed mutations, these computations allowed for comparisons across disease phases and vaccination status. Collectively, the virus substitution rate in non-vaccinated cattle was relatively high during early infection and decreased through disease progression. Contrastingly, in vaccinated cattle, the substitution rate was lower and relatively unchanged across progressing phases of disease. Although multiple instances of re-emergence of specific clades were found within the vaccinated animals, this phenomenon could not be inferred in the non-vaccinated cohort. Thus, viruses that transiently emerged in non-vaccinated animals did not re-occur at later time points. This superiority of clade-specific clearance in non-vaccinated animals may reflect differences in the host immune response during early [22] or late [28, 34] infection. One possible explanation is that a primed immune response in vaccinated animals led to fewer clades successfully seeding infection (i.e. stronger bottleneck).

It is likely that both evolutionary processes (clade emergence and within-clade evolution) are ultimately influenced by host immune factors acting upon the viruses’ intrinsic abilities to evade such pressures. In general, the higher proportion of substitutions, especially non-synonymous changes, present in structural as compared to non-structural genes (Table 1) suggests that conventional immunological pressures influence within-host virus evolution. These observations are consistent with the concept of molecular memory which suggests that the viral swarm retains previously adapted (e.g. escape mutant) viruses at low frequencies as an explanation for the observed complexity of FMDV populations [18, 58, 59].

One of the intentions of this study was to investigate viral genomic changes associated with different phases of infection defined as early, transitional, and carrier (persistent) phases. Consensus viruses sampled during early infection were notably diverse in their clade associations, which is of specific relevance as this is when infected animals are most contagious and likely to transmit disease [60, 61]. The vast majority of early consensus-level nucleotide changes corresponded to variants that were present at subconsensus level in the inoculum. This illustrates that the diversity present in the primary infecting virus population has a substantial effect upon the nature of the founding infection(s) suggesting that the role of immune-driven selection may be less influential than stochastic processes in the very early stages of FMDV pathogenesis.

The transitional phase of infection is the period during which animals either successfully clear the virus or develop into persistently infected carriers [28]. Therefore, this specific period is of particular interest due to the potentially significant selective pressures by the host immune response. Consequently, FMDV clade shifts occurred in every animal that was sampled through the transitional phase of infection. Recent investigations have suggested that virus clearance is associated with an activated cellular immune response [34, 42]. It is possible that transitional phase clade shifts are a result of this activated host response, and that virus populations unable to effectively evade this pressure are those that allow for complete clearance. Contrastingly, clade shifts were largely absent during persistent infection. This finding also may relate to the recent finding that cell-mediated killing as well as inflammatory- and apoptotic pathways are reduced in carriers [34, 42], potentially facilitating the stabilization of surviving virus populations.

The limited but clear emergence of amino acid substitutions within, and near, well-defined antigenic epitopes in the virus capsid inspired the analysis of capsid structural changes. Presumably, the strong selective pressure exerted by the adaptive immune response selects for epitope escape variants. Most of these substitutions occurred in different clade backgrounds which also suggests selective pressure dominating over stochastic drift. The only capsid substitutions that were detected earlier than 5 dpi were pre-existent variants that were present in the inoculum at high frequencies (nt positions 1903 (VP3 131) and 2615 (VP1 147) at 23.0% at 47.5%, respectively), suggesting some delay in mutation-evasion. Three of the amino acid substitutions that occurred within receptor-binding domains, namely VP1 144, VP1 199 and VP3 175 encoded qualitatively divergent amino acids, thereby drastically impacting local noncovalent bonding. This suggests that strong selection related to host cell entry may be driving these changes. The serine to arginine substitution at VP1 144 was found in all samples obtained later than 10 dpi from non-vaccinated animals, and in 2 out of 4 vaccinated animals that were followed through to the persistent phase of infection. Consequently, serine at VP1 144, which was present at a frequency of approximately 98% in the inoculum, was maintained through to the persistent phase of infection in only two vaccinated animals. This VP1 S144R change arose in the background of clades I-V by way of two different codon mutations, strongly suggesting that arginine is advantageous at this position. This is also consistent with most wild-type FMDV sequences, which overwhelmingly contain arginine at this position [62]. The dynamic appearance and disappearance virus populations coupled with substitutions at the major antigenic sites suggests the existence of an antigenic selection process during long-term (persistent) infection. Additional investigations aimed at providing a comprehensive analysis of FMDV structural adaptations are ongoing.

## Conclusions

This investigation was the first study to evaluate the full FMDV coding sequence across multiple animals through several distinct stages of infection in a natural host species. Characterization of consensus viruses in animal samples indicated dynamic patterns of emergence of pre-existing minority virus haplotypes (clade shifts). These clade shifts were responsible for the majority of consensus-level changes in multiple sample types. However, individual nucleotide substitutions, not otherwise associated with particular ancestral genotypes also occurred, presumably through point mutation. In every animal, the transitional phase of infection included a clade shift, whereas few clade shifts occurred during persistent infection, despite ongoing genomic changes. Overall, the data presented herein suggest that substantially different immunological, virological, and pathological selective processes occur during distinct phases of FMDV infection. More specifically, these findings suggest that the combination of host selective pressures and viral evolutionary mechanisms which enable the establishment of FMDV persistent infection may be distinct from the processes which allow maintenance of persistence.

## Acknowledgements

The authors would like to thank EJ Hartwig, GR Smoliga, and BP Brito for support in sample preparation and sequence finishing as well as WM Fischer for helpful insights. IF is a recipient of a Plum Island Animal Disease Center Research Participation program fellowship, administered by Oak Ridge Institute for Science and Education (ORISE) through an interagency agreement with the US Department of Energy.

